# Uncertainty Quantification in Acoustic Impedance of Atlantic Salmon Fish Scale using Scanning Acoustic Microscopy

**DOI:** 10.1101/2023.12.20.572508

**Authors:** Komal Agarwal, Shivam Ojha, Roy Ambil Dalmo, Tore Seternes, Amit Shelke, Frank Melandsø, Anowarul Habib

## Abstract

Scanning Acoustic Microscopy (SAM) emerges as a versatile label-free imaging technology with broad applications in biomedical imaging, non-destructive testing, and material research. This article presents a framework for the estimation of stochastic impedance through SAM, with a particular focus on its application to the salmon fish scale. The framework leverages uncertain reflectance, marking its pioneering application to uncertainty quantification in the acoustic impedance of fish scales through acoustic responses. The study uses maximal overlap discrete wavelet transform, to decompose acoustic responses effectively and is further processed to predict the acoustic impedance. To establish the effectiveness of the proposed framework, well-known materials like a pair of target medium (polyvinylidene fluoride) and reference medium (polyimide) are employed for impedance characterization. Results demonstrate over 90%accuracy in PVDF impedance estimation, validating the framework. A stochastic impedance map, using Kriging with a Gaussian variogram, offers insights into the complex biomechanics of a fish’s scale.

## 1 Introduction

Scanning acoustic microscopes (SAM) are widely used in materials science and biology for non-destructive and non-invasive imaging of surface and internal structures^1,2^. SAM enables the inspection of materials and can provide quantitative information like the thickness of the sample, the velocity of sound in the sample, subsurface defects, and Young’s modulus^3,4^. This versatile technology is widely used for mechanical characterization, including surface and subsurface evaluations, structural health monitoring (SHM) of composite structures, defects in polymer circuits, and analysis of anisotropic phonon propagation^5–7^. Many studies have scrutinized the acoustic characteristics of various biological tissues and biological-like tissues, with an increasing focus on evaluating frequencies surpassing 25 MHz^8,9^. This heightened attention is attributed to the rising significance of high-frequency ultrasound in life science applications. SAM’s non-destructive, non-invasive, and deep penetrating imaging capabilities also unlock the potential of acoustic imaging in biomimetics. SAM technology offers comprehensive evaluations of both biological samples and bio-mimicked synthetic materials, providing invaluable insights for the advancement of material development. SAM has been previously used for imaging details of bone and dental structures^10–12^.

In the field of biomimetics, researchers worldwide draw inspiration from the natural world to create materials that fulfill crucial functions and address intricate challenges^13–15^. Among these natural materials, fish scales, especially those of the salmon (*Salmo salar*), have garnered significant attention due to their extraordinary flexibility, strength, resistance to penetration, and lightweight properties, influencing the dynamic motion of fish. The unique attributes of salmon scales have positioned them as a focal point in biomimetic armor designs, where researchers seek to replicate and leverage their advantageous features.

Beyond the realm of biomimetics, these scales play a crucial role in the physical protection of fish. Furthermore, for more than a century, salmon scales have been indispensable for estimating fish age and growth^16^. Considering that salmon populations are confronted with threats like habitat degradation, pollution, diseases, and over-exploitation^17^, understanding the age distribution within fish populations becomes vital for effective fisheries and conservation management^18,19^. The multifaceted potential of salmon scales in bio-inspired armor and disease diagnosis underscores their paramount importance. Traditional optical observation methods encounter challenges such as limited depth of penetration and difficulties in imaging live samples^20^. Additionally, the use of lasers in conventional methods poses a potential risk of damaging sensitive biological samples^21^. Exploiting the advanced capabilities of SAM, it emerges as a particularly suitable approach for obtaining detailed images of fish scales. This not only enhances our comprehension of fish biology but also contributes to advancements in aquatic ecology and biomimetics. SAM’s ability to overcome the limitations of traditional observation methods positions it as a valuable tool for studying biological structures with precision and depth.

The acoustic response obtained after SAM has a variety of frequency bands that arise due to the multiple reflection interfaces and these frequencies are evolving with time which ultimately creates the need for time-frequency analysis of decomposition. The basic methods for observing such signals are short-term Fourier transform (STFT)^22^ and analysis of such signals are time windowing and Fourier transform, which breaks signals into temporal and frequency components^23^. Time windowing is difficult as it requires precision in the time selection as well and multiple signal bands can overlap within the time window causing issues in decomposition. Moreover, the Fourier analysis involves pre-processing steps like data windowing. Advanced tools like Wavelet transform have better signal decomposition in the multi-scale resolution^23^. In wavelet decomposition, discrete wavelet transform (DWT) imposes restrictions on the length of signals, which should be the multiple of powers of two. This limitation restricts the DWT application, and decomposition depends on whether the event span falls within a wavelet averaging window or not^24,25^. Thus, this work utilizes the maximal overlap discrete wavelet transform (MODWT) which persists down-sampled values at each decomposition level^23,25^. Thus, attributing the advantages of the MODWT, its combination with other methods will create an effective and efficient approach for extracting the characteristic features through the acoustic response of fish scales.

In this work, we determine the acoustic impedance of the fish scales. SAM utilizes focused acoustic signals, emerging as a powerful tool for tissue characterization and elastic parameter imaging^26–29^. This article introduces acoustic impedance microscopy of fish scales as a non-invasive method, aiming to image local acoustic impedance distribution which is a parameter closely related to sound speed and potentially valuable for tissue characterization. Moreover, this article presents the pioneering application of the uncertainty in the reflectance for estimating the stochastic impedance of the fish scale. By exploiting the relationship between acoustic impedance, sound speed, and density, this methodology enables micro-scale imaging through acoustic response.

The article presents the process of extracting dominant frequencies from the acoustic signal through MODWT combined with a bandpass filter, thus building up a basis for the estimation of impedance. An uncertain reflection coefficient is considered to estimate the stochastic acoustic impedance. Lastly, Kriging is employed for spatial interpolation of impedance which uses both linear and Gaussian variograms. The Gaussian-based Kriging is also referred to as Gaussian process regression (GPR). It is flexible and can incorporate the uncertainty in the observed data while performing regression and handles complexities in non-linear mapping^30^. The algorithm is validated on known materials before application to bio-samples. Figure 1 depicts the overall strategy used in this article. Overall, this pioneering approach aims to deepen our understanding of fish biomechanics, offering improved insights into the broader marine ecosystem and revolutionizing the diagnosis of diseases and structural changes within marine communities.

**Figure 1.**
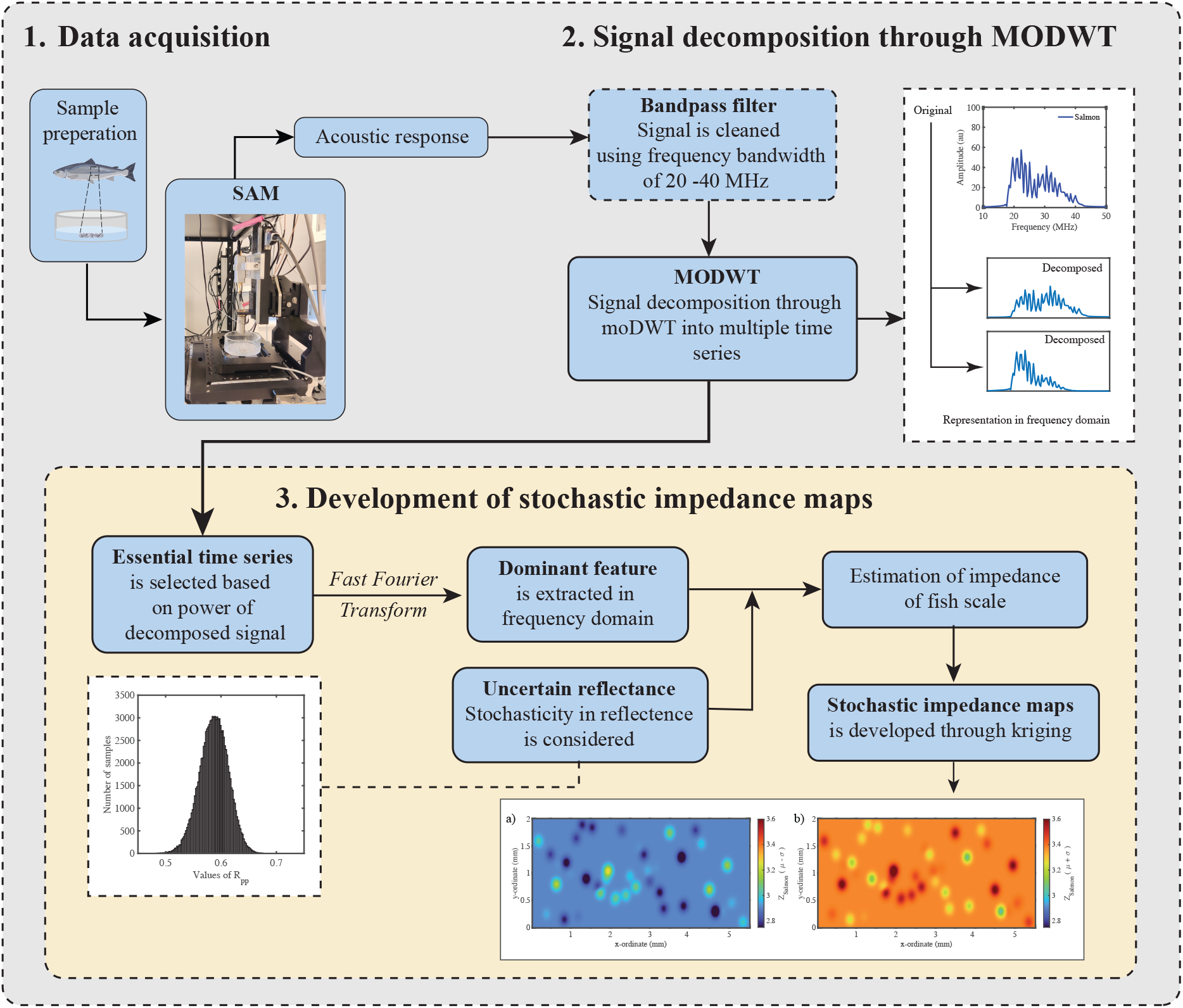
The schematic diagram depicts the entire process for creating the stochastic impedance map of the fish scale. The initial step involves SAM imaging, which records the images of multiple samples. In the second section, we decompose the acoustic response, and then we proceed with the decomposed time series. The final section illustrates the selection of essential time series and the development of stochastic impedance maps using uncertain reflectance.

## 2 Experimental procedure

### 2.1 Sample preparation

Salmon fish scales were obtained from healthy salmon obtained from the Tromsø Aquaculture Station. No ethical approval had to be obtained since the fish received no treatment before euthanization. The fish scales were carefully pulled from the fish skin, by gentle pulling with a plastic tweezer. This approach was adopted to ensure that the fish scale samples remained intact and undamaged during the extraction process.

A solution was prepared by dissolving 2 wt% of agarose (specifically, ultra-low gelling temperature agarose from Sigma Aldrich) in 10 ml of distilled water. To ensure proper mixing, we used a magnetic stirrer to agitate the mixture in a beaker, maintaining a temperature of 100 °C for 10 minutes. Afterward, we poured a thick gel into a Petri dish, and on top of this gel, we carefully placed a fish scale and gently pressed it into the agarose layer (Figure 2c). A schematic illustration of the sample preparation process is depicted in the following Figure 2. We gently poured cold distilled water into the Petri dish and promptly began the data collection process to preserve the sample’s integrity. The utilization of agarose was twofold in its purpose. Firstly, it was employed to preserve the freshness of the sample, ensuring that it remained in an optimal condition for imaging. Secondly, agarose acted as a reliable means to anchor the scale securely within the water bath, preventing any unwanted movement or displacement during the scanning process^31^. Additionally, agarose possesses an acoustic impedance that closely matches with water^32^.

**Figure 2.**
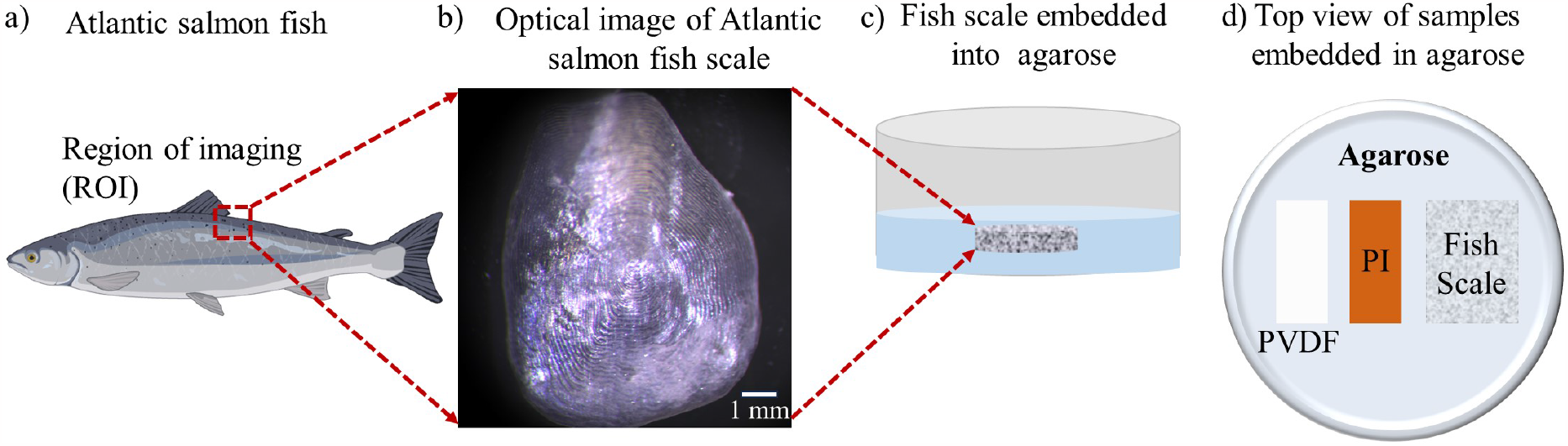
A schematic representation of steps involved in sample preparation. (a) shows the sampling ROI located near the dorsal fin of the Atlantic salmon used for SAM imaging, (b) shows an optical image of a fish scale taken from the ROI, (c) shows how an extracted fish scale is embedded into the agarose gel (blue), water is filled on top (gray) for coupling in SAM imaging, (d) shows the top view to illustrate the spatial arrangement of the reference samples PVDF and polyimide (PI) with respect to the target sample fish scale

To determine the acoustic impedance of the fish scale, two additional samples with known impedances, polyimide (PI) and polyvinylidene fluoride (PVDF), were also incorporated into the agarose gel, as illustrated in Figure 2d. This method enabled the acquisition of images for all the samples in a single session, simplifying the comparative analysis.

### 2.2 Scanning acoustic microscopic imaging

Figure 3 demonstrates a labeled representation of a SAM, utilized for capturing images of samples. SAM utilizes both reflection and transmission modes, each providing distinct insights into different aspects of the sample’s characteristics. The annotated image of SAM in Figure 3 highlights its key components or operational settings, serving as a reference for image acquisition. Further details regarding the operational principles for these modes can be found in the following literature^33,34^.

**Figure 3.**
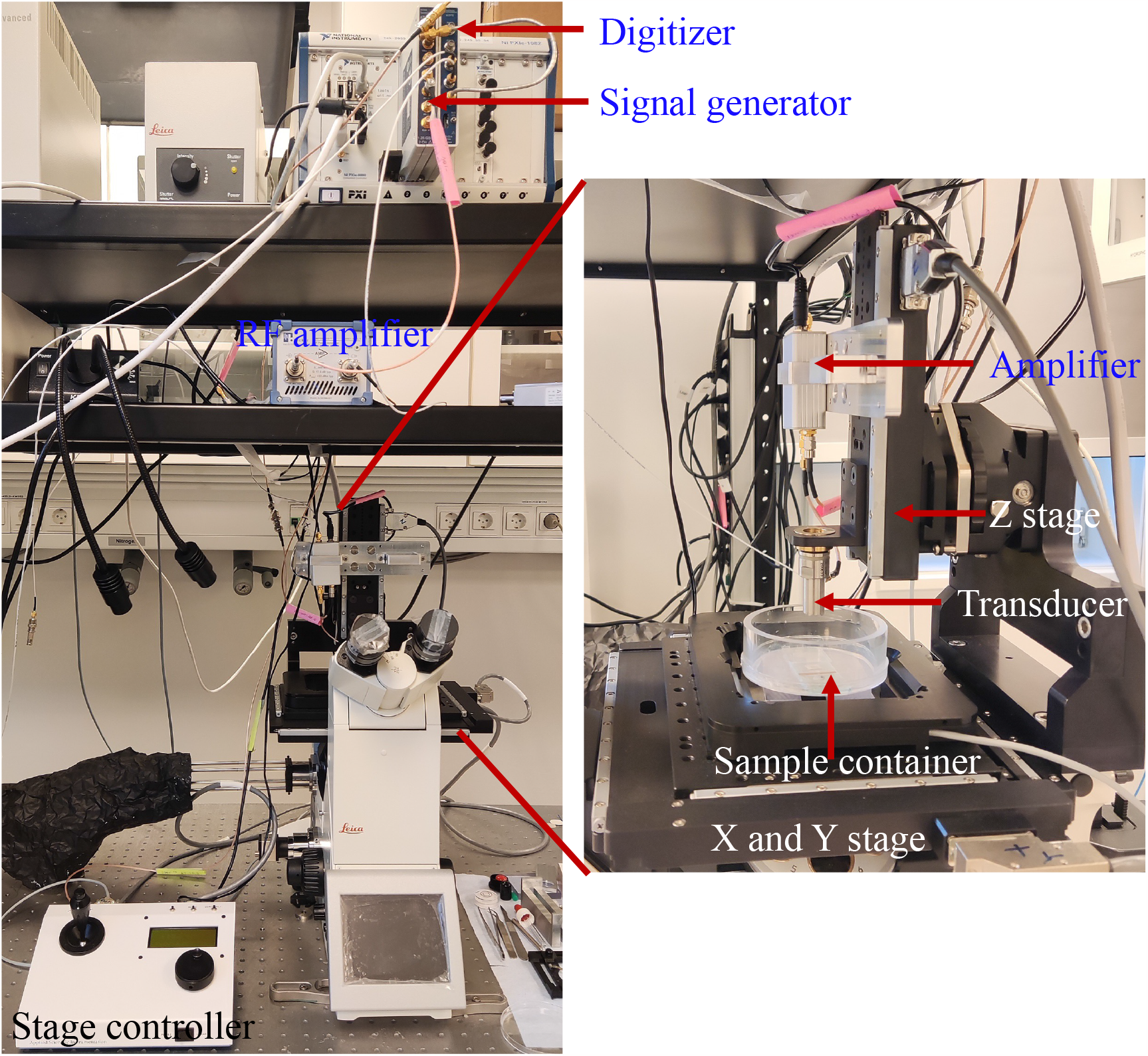
The figure shows a labeled image of the SAM that was used for image acquisition of samples in this article. The experimental setup displayed in the figure demonstrates all the essential components of the SAM.

In this article, we primarily focus on using the reflection mode for scanning samples. To accomplish this, a frequently used method involves employing a concave spherical sapphire lens rod to concentrate acoustic energy through a coupling medium, often water. Ultrasonic signals, which are excited by a signal generator, are directed toward the sample. When these signals bounce back from the surface of the sample, the reflected waves are captured and converted into a digital signal, typically known as an A-scan or amplitude scan.

To generate a C-scan image of the sample, this process is performed at several different positions within the XY plane. Another perspective on visualizing a C-scan is to regard it as the integration of A-scans in two dimensions. We developed a custom-designed ultrasound scanning system by combining a Leica DMi8 inverted microscope with a high-precision ASI MS-2000 XYZ scanning stage. LabVIEW software was utilized to control both the scanning stage and other microscope components. The ultrasound functionality was realized through the integration of PXIe FPGA modules and FlexRIO hardware from National Instruments, which were installed in a PXIe chassis (PXIe-1082). The hardware consisted of an arbitrary waveform generator (AT-1212) and a 3W RF-amplifier^35^ for ultrasound pulse generation. Additionally, a 12-bit high-speed (1.6Gs/s) digitizer (NI-5772) was used for reflected signal recording.

We used a 30 MHz PVDF-focused Olympus transducer with specific dimensions - a 6.35 mm aperture and a 12 mm focal length^35^. To ensure accuracy, the thickness of the PVDF, PI film, and fish scale was measured using a digital micrometer, with approximate thicknesses of 105 *μm*, 130 *μm*, and 100 *μm*, respectively. We scanned the samples within a scanning area of 10 mm *×* 2 mm, and each pixel represented a size of 50 *μm* in both the x and y dimensions. We performed scans that covered a range from 200 *μm* above to 800 *μm* below the focal point of the thickest sample (PI), using a step size of 20 *μm*. This meticulous approach was taken to guarantee that all samples, regardless of their varying thicknesses, were imaged at their focal planes. Additionally, it allowed us to obtain z-scans of the samples at various depths, ensuring comprehensive data acquisition.

Additionally, to view the entire features of a fish scale, we imaged a full/complete area of a fish scale using SAM. Figure 4 shows the acoustic amplitude image of the full fish scale captured at the focal point plane, with a scanning area of 13 mm *×* 13 mm, and with x and y pixel size of 50*μm*. The circular ridges that are present on top of a fish scale (as also seen in the optical image of Figure 2d) can be seen clearly in this amplitude image.

**Figure 4.**
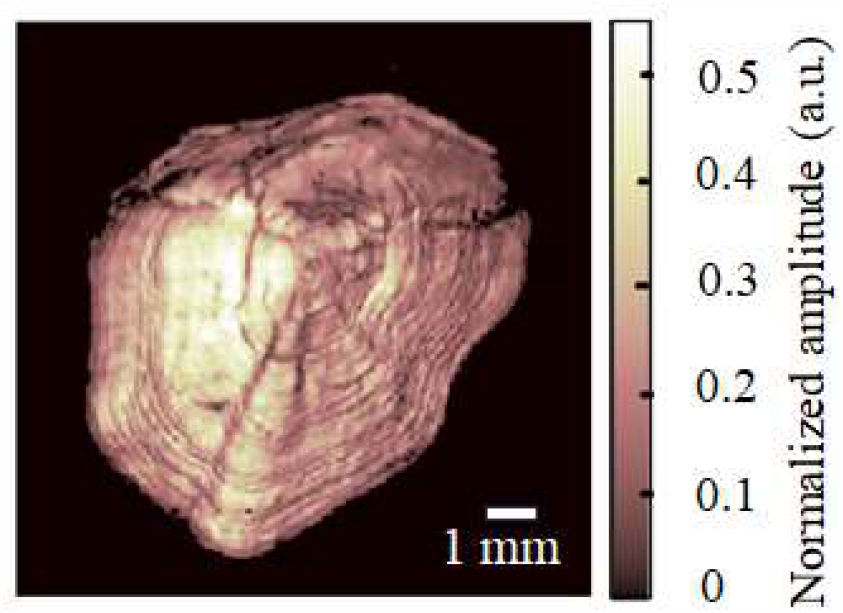
Scanning acoustic microscopic image of the salmon scale. Imaging was performed employing a 30 MHz focused polymer transducer. The scanning area was 13 mm *×* 13 mm and the step size was 50 *μ*m in both directions.

## 3 Theory of acoustic waves

Acoustic microscopy functions in a non-destructive manner, employing the penetration of acoustic waves to render internal features visible. This propagation of acoustic waves induces both normal and shear stress within the medium, playing a crucial role in categorizing the waves into P waves (Primary or Pressure waves) and S waves (Secondary or Shear waves). In the context of a planar interface, the incidence of the P-wave results in four distinct wave components: a) reflected P wave (RPP), b) reflected S wave (RPS), c) transmitted P wave (TPP), and d) transmitted S wave (TPS). To express these wave phenomena mathematically, the wave potentials can be articulated as follows^36^:

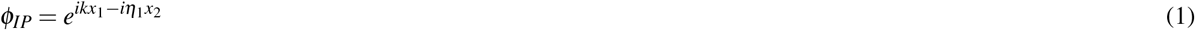

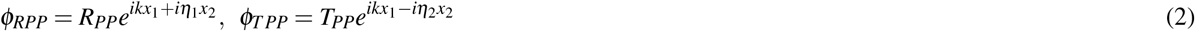

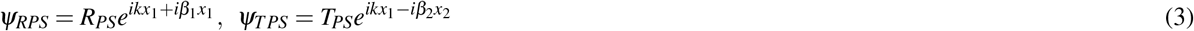

where subscript numbers 1 and 2 represent the medium and,

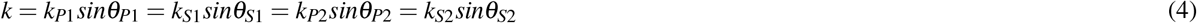

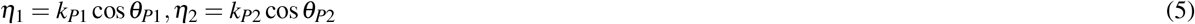

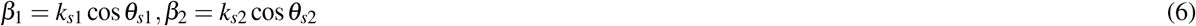

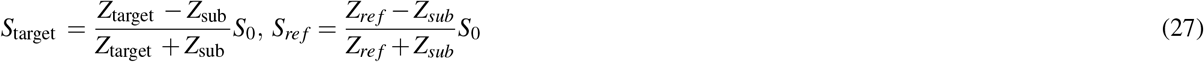

Where *k*_*P*1_, *k*_*P*2_, *k*_*S*1_, *k*_*S*2_ represent the wave numbers of P and S waves traveling in materials 1 and 2, respectively, and other symbols carry their standard meanings. In the specific scenario of normal incidence of the P-wave, i.e., *θ*_*P*1_ = 0, *θ*_*P*2_ = 0, *θ*_*S*1_ = 0, *θ*_*S*2_ = 0, the continuity of displacements and stress conditions at the interface yields the following simplified matrix:

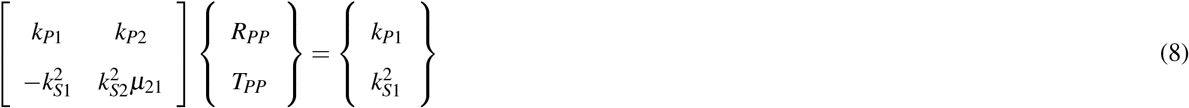

Here, *μ*_21_ = *μ*_2_*/μ*_1_ represents the ratio of lame’s constants, and the values of reflectance *R*_*PP*_ are determined as follows:

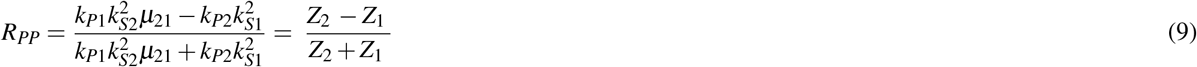

where, *Z*_1_ and *Z*_2_ denote the acoustic impedance of the medium 1 and 2, respectively.

### 3.1 Stochastic formulation of reflectance

In the current work, it is assumed that wave number is not deterministic; rather, it follows the normal distribution with some mean and standard deviation. An uncertain wave number or momentum uncertainty implies a spread or uncertainty in the wave function’s spatial properties. It suggests that the wave particle’s position is not precisely defined but rather distributed over a range of positions. Moreover, from the signal perspective, there can be closely spaced frequencies arising due to momentum uncertainty in the acoustic signals.

Let us consider the wave number to be normal distributed, this implies *k*_*P*1_ *∽ N* (*μ*_*P*1_, *σ*_*P*1_), *k*_*S*1_ *∽* (*μ*_*S*1_, *σ*_*S*1_), and *k*_*P*2_ *∽ N* (*μ*_*P*2_, *σ*_*P*2_), *k*_*S*2_ *∽ N* (*μ*_*S*2_, *σ*_*S*2_), then mean and variance of the reflectance will be derived using logarithmic transformation. Taking log on both sides of Eq. 9, the resulting expression would be:

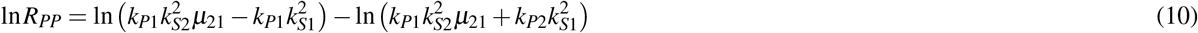

By expanding the right side of the equation using Taylor series expansion of ln(*x*) up to second order approximation as ln 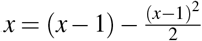, we obtain:

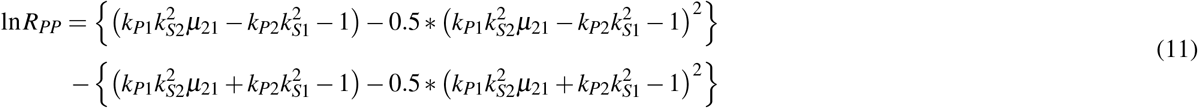

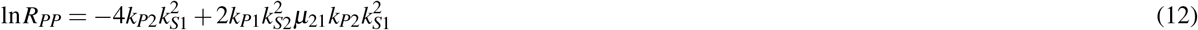

Thus, we have obtained an approximation of ln *R*_*PP*_. The derivation for the mean and variance are discussed in subsequent subsections.

### Mean of ln **R**_**PP**_

Through operating expectation on both sides of Eq. 12, and assuming all the wave numbers to be independent, the mean of the ln *R*_*pp*_ is written as:

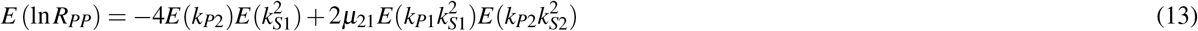

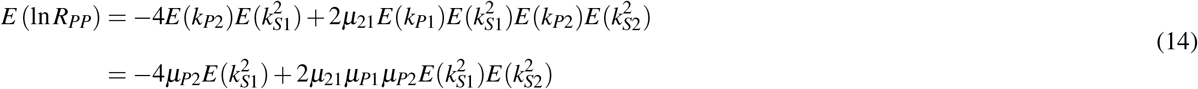

For the evaluation of higher moments i.e., *E*(*X* ^2^), *E*(*X* ^4^), we have used the properties of the moment generating function.

Thus, if *X* follows the normal distribution, then, its moment generating function is written as 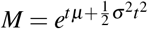, Therefore, the moments can be written as

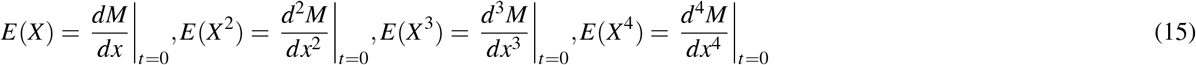

Solving the above equation, we can write:

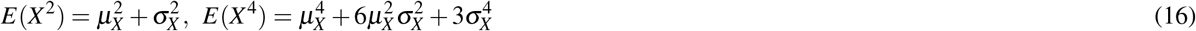

Hence, the mean can be written as follows:

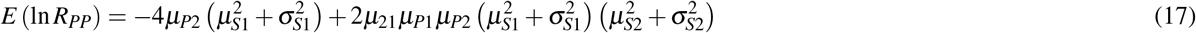

### Variance of ln **R**_**PP**_

Through operating variance on both sides of Eq. 12, and assuming all the wave numbers to be independent, the mean of the ln *R*_*pp*_ is written as:

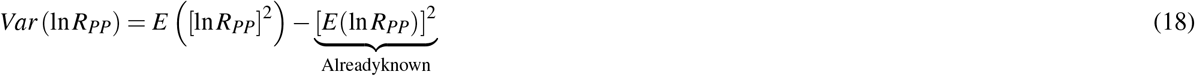

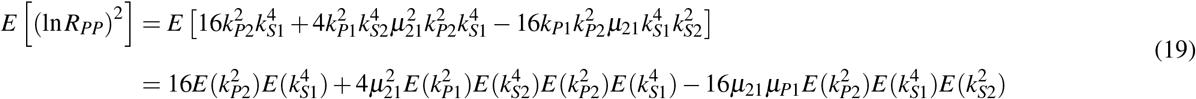

Where 2^th^ and 4^th^ moments are calculated using the moment regenerating function as follows:

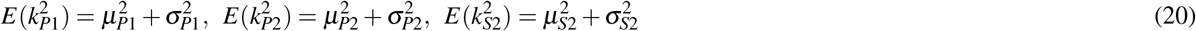

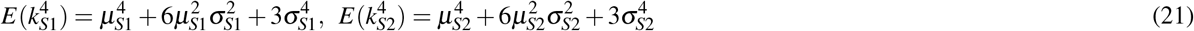

### Evaluating mean and variance of R_PP_

Let mean of the *R*_*PP*_ be *μ*_*R*_, then using Jensen’s inequality, the approximate values of *E*(*R*_*PP*_) is given by:

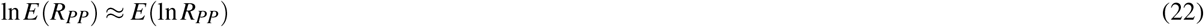

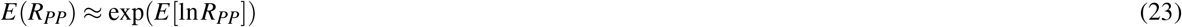

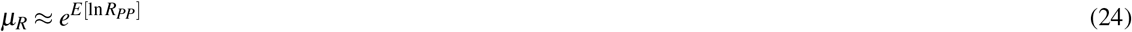

Similarly, the variance of the *R*_*PP*_ is approximated by using delta method:

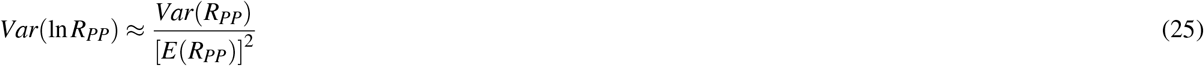

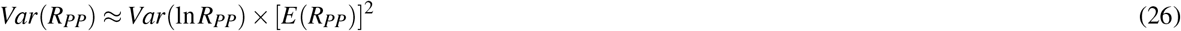

### Estimation of Impedance

Considering the characterization frequency of the transmitted/incident signal, the reflected signal through reference and target medium be *S*_0_, *S*_*re f*_ and *S*_*target*_ respectively. Then, we can write the following relations^36–38^:

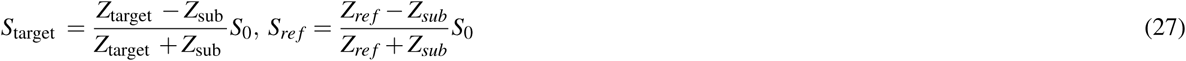

where *Z*_*target*_, *Z*_*re f*_, and *Z*_*sub*_ represent the acoustic impedance of the target, reference, and substrate, respectively. The measurement is possible only for the *S*_*re f*_ and *S*_*target*_ and *S*_0_ cannot be measured directly. Let 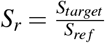, and 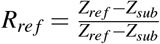 The acoustic impedance of the target medium is subsequently is given by:

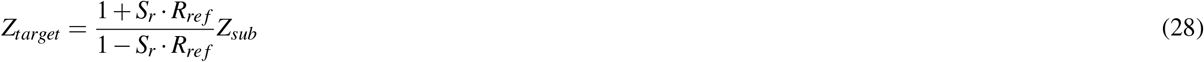

Considering the uncertainty in the *R*_*re f*_ with mean *μ*_*R*_ and standard deviation *σ*_*R*_. For the calculation of *μ*_*R*_ and *σ*_*R*_, appropriate wave numbers are selected with proper mean and variance. Now, The values of estimated mean *μ*_*Z*_ and bounds *μ*_*Z*_ *±σ*_*Z*_ for the target impedance are given by:

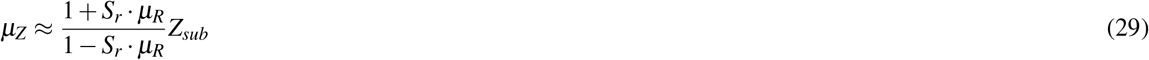

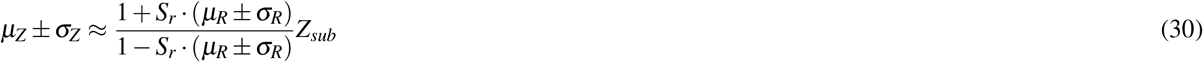

Thus, the current work considers uncertainty in reflectance while estimating the acoustic impedance of the target medium. Moreover, it is important to note that the accuracy of the approximation increases if fourth-order logarithmic expansion is considered instead of second-order expansion.

## 4 Methodology for predicting stochastic impedance maps

In the current work, the stochastic impedance map of the fish scale is developed through acoustic microscopy using the maximal overlap discrete wavelet transform (MODWT) and uncertainty in reflectance. The proposed methodology assumes variation in the horizontal plane only and assumes the homogeneous medium in the vertical plane, thereby, avoiding the cause of multiple refections from the same medium. Moreover, due to various interfaces apart from a homogeneous medium, the reflected response of the acoustic imaging contains multiple peaks having different frequency components that are transmitted and reflected through the various interfaces resulting in the formulation of a time-frequency analysis problem. This response is first passed through the bandpass filter to remove unwanted noise and then MODWT is applied to decompose the signal into multiple time series having its own wavelet and scaling coefficients for different levels.

### 4.1 Maximal overlap discrete wavelet transform (MODWT)

In the discrete wavelet transform (DWT), the signal is decomposed into different component levels through filtering and down-sampling, where decomposition at each level is characterized through its approximation and detailed coefficients^23–25^. Further, Mallat’s algorithm is used for the practical application of DWT which contains a pair of filters: a low-pass filter (scaling function) and a high-pass filter (wavelet function). The application of Mallat’s algorithm continues recursively on the resulting low-pass signal till the required decomposition level. For a discrete signal, **X** = *{X*_*t*_, *t* = 0, 1, *…, N −* 1*}*, the DWT computes the wavelet coefficient for the discrete wavelet of scale 2 ^*j*^ and location 2 ^*j*^*k* using the following equation:

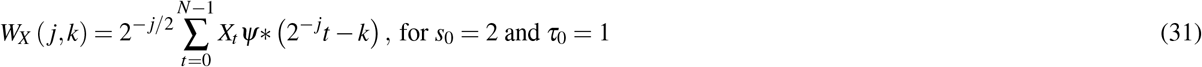

where *W*_*X*_ ( *j, k*) is the wavelet coefficient and *N* is an integer power of two.

However, DWT is incapable of handling shift-invariance and has narrow frequency resolution, thus restricting its application to the fixed signal length with an integer multiple of a power of two^23^. A modified version called MODWT divides the frequency band of the input signal into scaling and wavelet coefficients using low- and high-pass filters, that is, scaling and wavelet filters. MODWT can be properly defined for arbitrary signal length and it achieves redundancy through an oversampled representation which enables more accurate statistical analysis^25^.

Let *X* = *X*_*t*_, *t* = 0, …, *T −* 1, be the time series data, then, the j^th^ level wavelet and scaling filters are denoted as 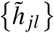 and 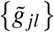, respectively. The scaling and wavelet coefficients calculated by Mallat’s algorithm are described as follows:

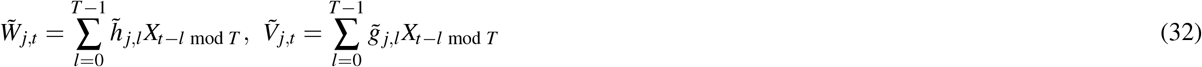

where *j* = 1, 2, …, *J*_0_ is the level of wavelet decomposition, and ‘mod’ denotes the remainder of dividing two numbers. The wavelet 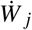 and scaling 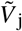 coefficients vectors of MODWT are written as:

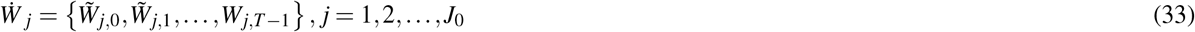

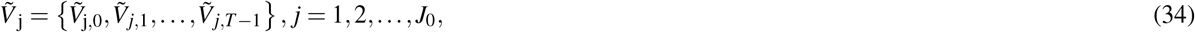

where 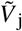 and 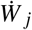 are related to the smallest and highest frequency components of the original signal 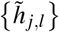, and 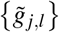 are the j^th^ level MODWT high-pass filter 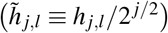 and low-pass filter 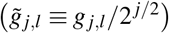 and *J*_0_ is the highest decomposition level. For the level 3 decomposition, the MODWT decomposes an original signal *X* into a low-pass filtered approximation component (A_3_) and high-pass filtered detail components (*D*_1_, *D*_2_ and *D*_3_). The equations of MODWT-based multi-resolution analysis can be used for the reconstruction of decomposed signal and are written as follows:

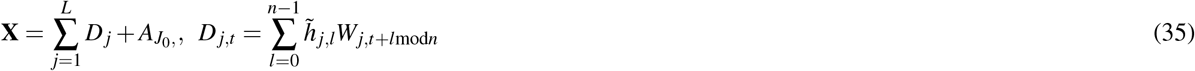

Through using the above equations, the signal is reconstructed for each level of decomposition. The next challenge is to select the appropriate decomposed time series for further processing which is discussed in subsequent sections.

### 4.2 Selection of essential time series and estimation of impedance

Once, the signal is decomposed into wavelet and scaling coefficients, it is important to reconstruct the decomposed time series and select the essential time series. In order to reconstruct the decomposed time series, the necessary decomposition levels are selected. The wavelet coefficients associated with those levels were passed through inverse wavelet transform (iMODWT) to obtain the decomposed time series. Next, the power of each decomposed time series is calculated and normalized with the maximum power. Thus the time series having maximum power relative to each other is selected as the essential time series. Thus, this essential time series is then transformed into the frequency domain to extract the character’s tic feature of the response. In the frequency domain, the frequency corresponding to the maximum amplitude is assumed to be the signal-characterizing feature and is used in the calculation of acoustic impedance.

Using the uncertain reflectance, the estimated impedance has its mean value as well as *μ* + *σ* and *μ −σ* values, thus giving rise to the stochastic prediction of impedance. In the current work, it is referred to as stochastic impedance. The impedance is first calculated at sufficient points, sampled using Latin hypercube sampling, spread over the whole domain, and then kriging is performed for all the three values of stochastic impedance i.e., *μ, μ* + *σ*, and *μ −σ* to find the impedance over the complete domain and stochastic impedance maps are generated to analyze the surface of fish scale. The basic understanding of the kriging is presented here.

Kriging^39^ is a method of interpolation based on the Gaussian process. It develops a meta-model of a partially observed function (of time and/or space) with an assumption that this function is a realization of a Gaussian process (GP)^30,40^, and thus the regression is also referred to as the Gaussian process regression (GPR). In GPR, the kernel functions are assumed, and the hyperparameters of the kernel functions are computed from observation via maximizing the log-marginal likelihood function. In the aspects of Kriging, the kernel function can be interpreted as a variogram, and two types of variogram are used in the current work, i.e., linear and Gaussian. Let the known locations be represented by 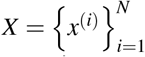 where *x* is the two-dimensional vector and the observed state at these locations is given by *y* = {*y*^(1)^, *y*^(2)^, *…, y*^(*N*)}^ . Let the meta-model between the observed state and known locations is given by:

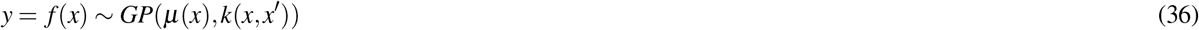

where *μ*(*x*) is the mean function and *k*(*x, x*^*′*^) is the kernel or covariance function. By definition, the prior of the observed state vector is Gaussian and is given by *p* ( *f* (*x*) | *x*) = *N* ( *f* (*x*) | *μ, K*). Given the prediction locations as *x*_*∗*_, the predicted state *y*_*∗*_ = *f* (*x*_*∗*_) is also jointly Gaussian and is given by

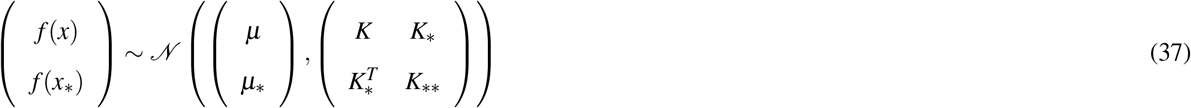

Using Bayesian transformation, the posterior mean and variance of the predicted state are written as

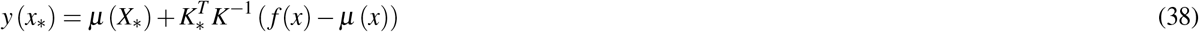

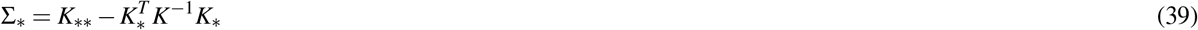

Lastly, the proposed methodology can be summarised as follows:

- Removal of noise through a band-pass filter
- Filtering of cleaned signal through MODWT
- Selection of essential time series
- Picking up the dominant frequency
- Estimation of mean impedance and its bounds
- Development of stochastic impedance maps

## 5 Results and Discussion

### 5.1 Validation of the proposed framework through known material

The accuracy of the proposed framework has been rigorously tested considering polyvinylidene fluoride (PVDF) as the primary material of interest, utilizing polyimide (PI) as the reference sample. The experimental process, detailed in Section 2, was meticulously described, recording the acoustic response of the system under study. In the experiment, the response has been obtained at nodes whose X and Y locations are represented by index i.e., Index (i) and Index (j). However, for brevity purposes, the results of only a particular node are presented. The response signal encompasses a spectrum of frequencies that evolve over time or we can say the same frequency is present at multiple time points. Thus, to separate the essential characteristics of the signal, the signal is first passed through the bandpass filter, and then decomposition is carried out using MODWT. This decomposition leads to the various time series at different decomposition levels represented by (1), (2), (3), (4) which are shown in Figure 5.

**Figure 5.**
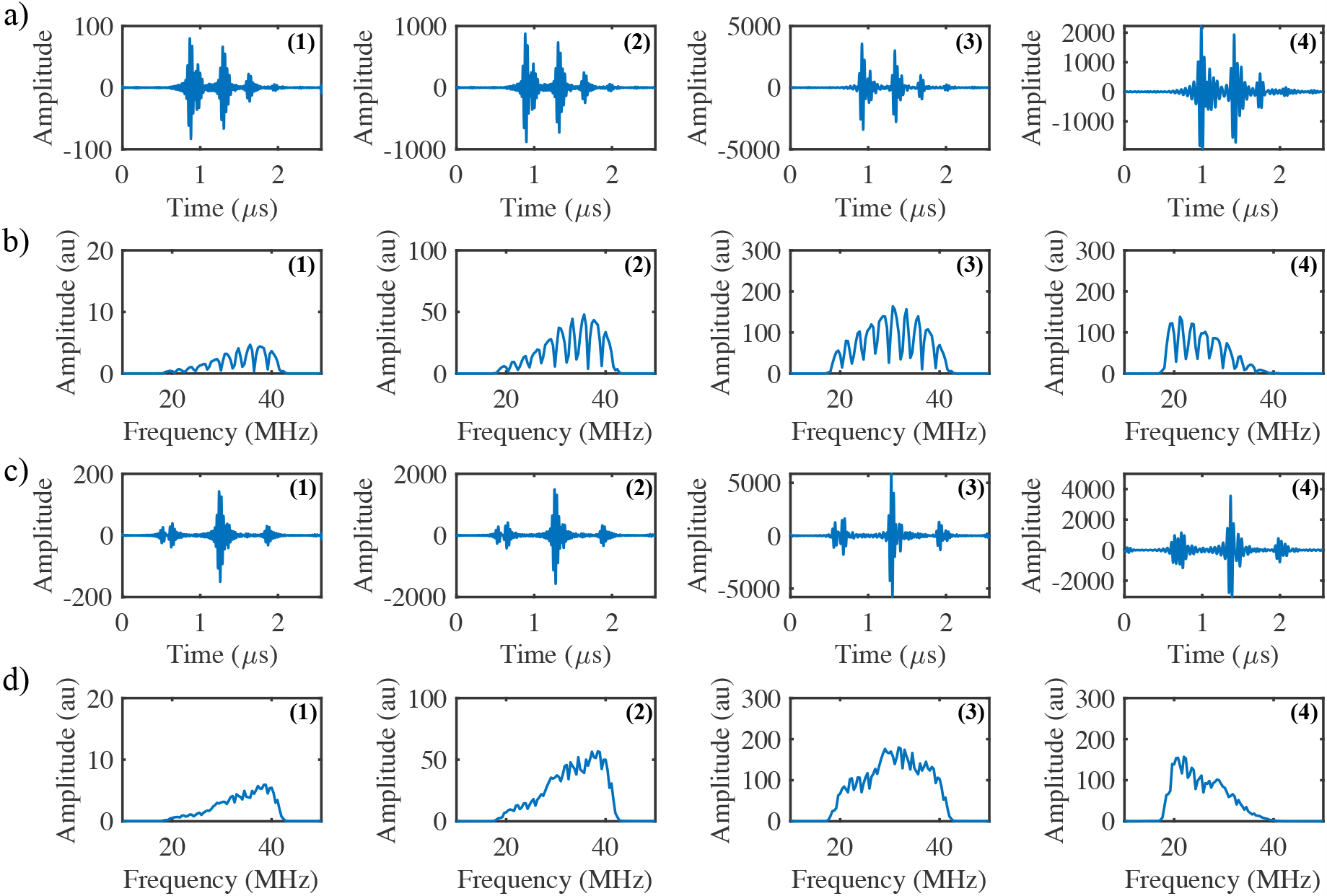
It demonstrates the decomposition of the acoustic response signal into multiple decomposed time series after MODWT in both the time-domain (a), (c) as well as frequency domain (b), (d). Figure (a) and (b) highlights the characteristics of the response signals PVDF at different decomposition level and figure (c) and (d) highlights the characteristics of the response signal of polyimide (PI) at various decomposition level.

While reconstructing the decomposed time series from wavelet and scaling coefficients of necessary level as described earlier, the coefficients associated with those levels were passed through inverse wavelet transform (iMODWT), and the power of each decomposed time series is calculated and normalized with the maximum power. The time series that has maximum energy content is selected as the essential time series and considered as the filtered signal. Figure 6 shows the normalized power of each decomposed time series with respect to others.

**Figure 6.**
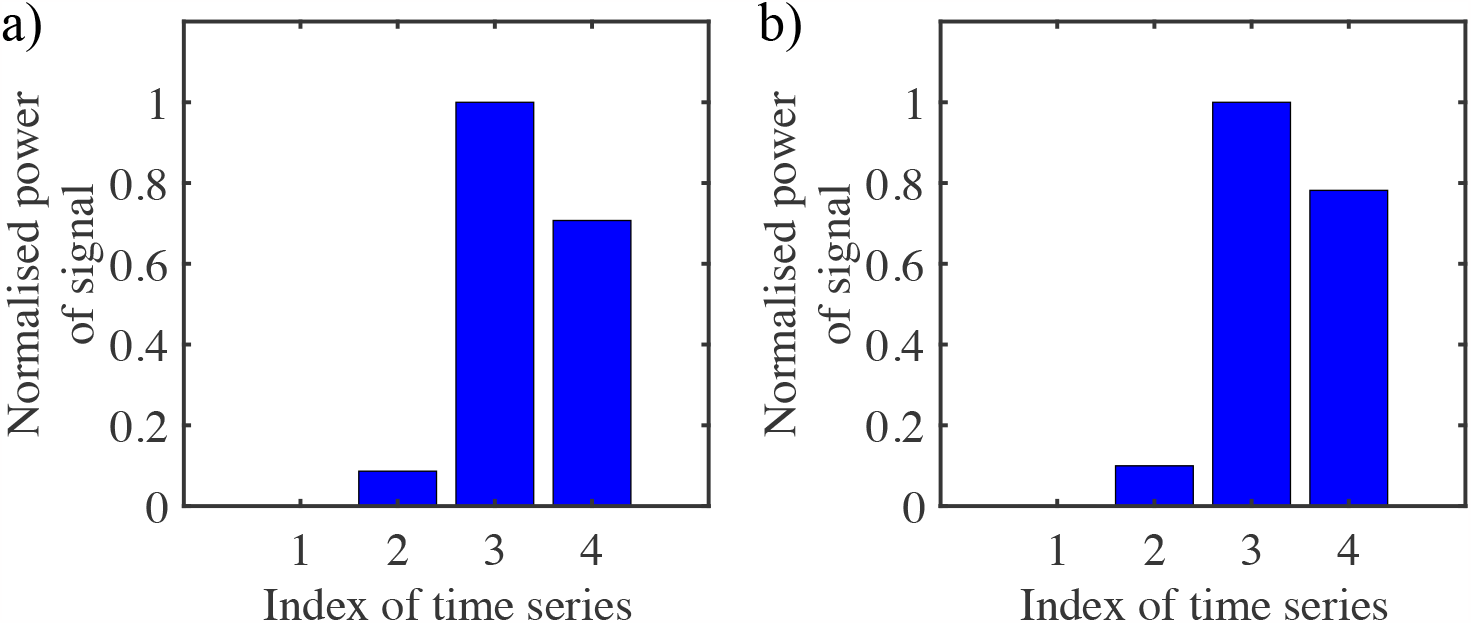
The figures represent the normalized power of each decomposed time series with respect to others for (a) PVDF and (b) polyimide (PI) samples, providing the basis for the selection of the dominant time series.

The essential characteristics of the signal that are directly correlated to the properties of the specimen are extracted from this filtered response. However, the changes in the time domain are difficult to interpret and therefore the responses are transformed to the frequency domain where the predominant peak frequency which corresponds to the maximum amplitude is selected as a representative of the specimen characteristics as presented in Figure 7.

**Figure 7.**
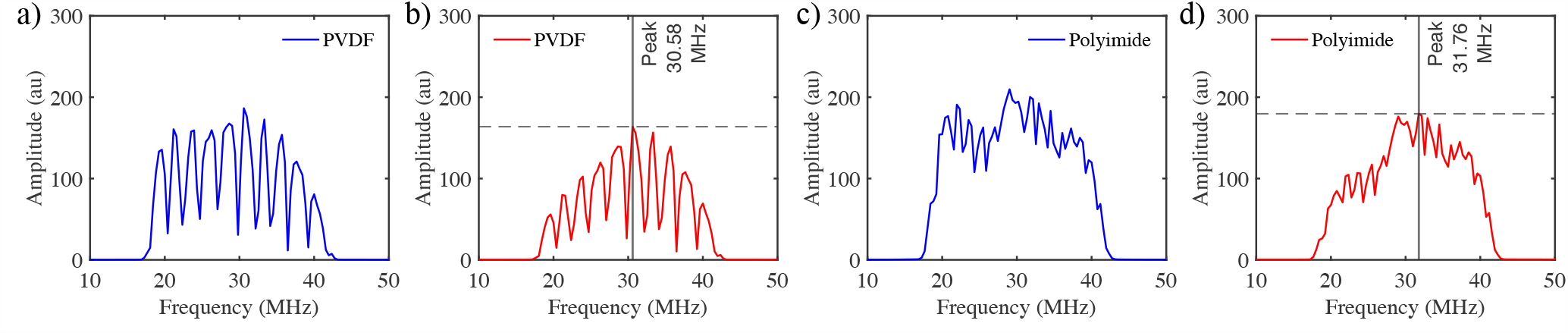
Displayed in the figures are the frequency spectra of the true responses (a) and (c), as well as the wavelet-transformed signals (b) and (d). These representations provide the identification of primary frequencies.

Figure 7(a, c) illustrates the frequency spectra of the true responses of the reference sample PVDF and target medium polyimide (PI). These obtained frequencies will provide the *S*_*ratio*_ as described in the theory section. The accuracy of the complete algorithm is determined through accurate estimation of these frequencies because the *S*_*ratio*_ is found to be very sensitive in the calculation of impedance. The frequency spectra of the filtered or wavelet transformed signal are shown in Figure 7 (b and d). The dominant peak from the frequency spectra of the filtered signal is obtained and shown in Figure 7 (b and d). After extraction of dominant frequencies, the stochastic impedance of the PVDF is calculated and thus, the estimated mean value is compared with the true impedance value of the PVDF along with the predicted *μ* + *σ* and *μ −σ* . The uncertainty in the reflectance which is considered in the current work is presented in Figure 8 (a).

**Figure 8.**
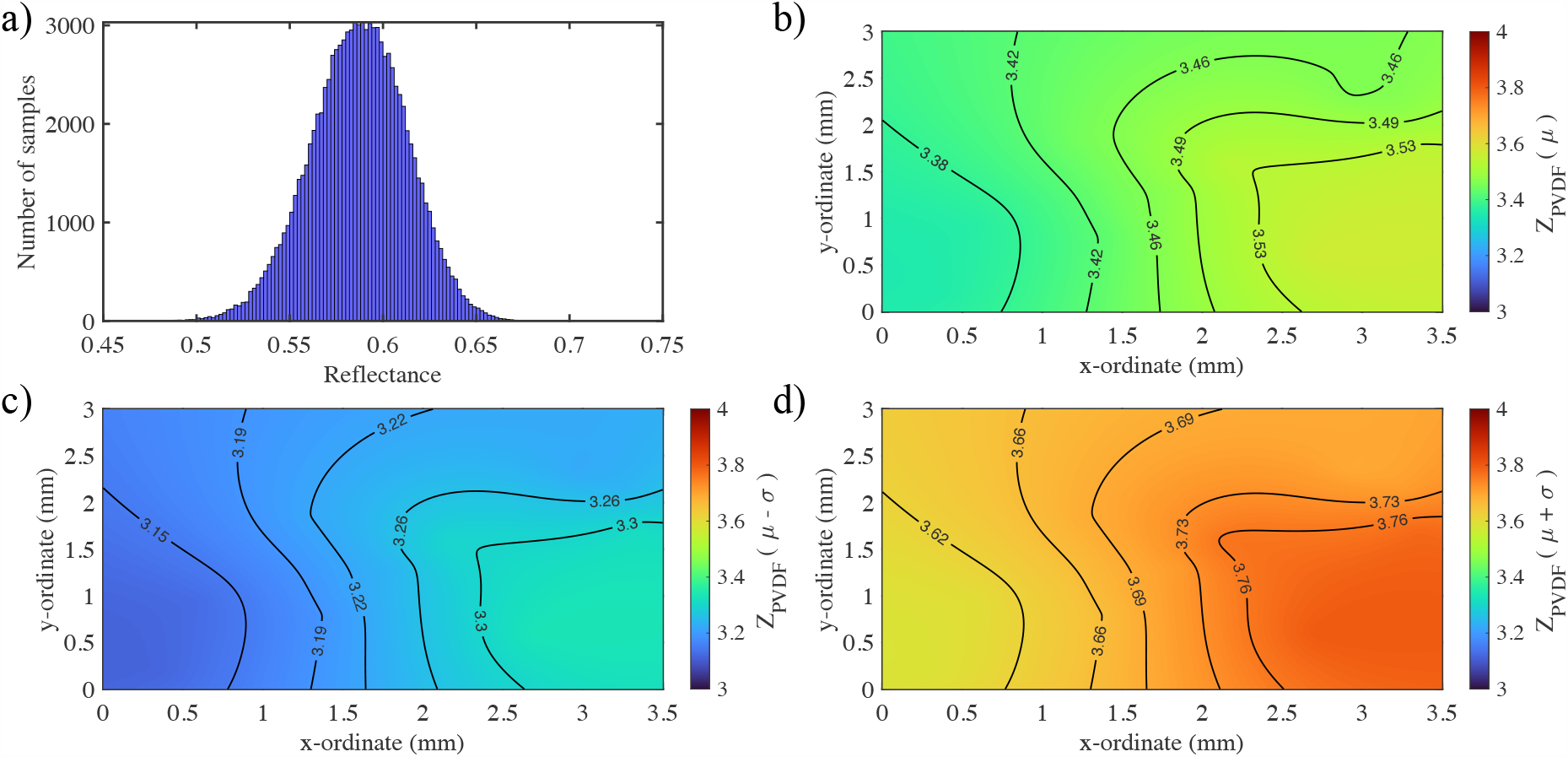
(a) Illustrates the considered variation in the reflectance (b) illustrates the mean value of the impedance (c) and (d) demonstrates the estimated *μ −σ* and *μ* + *σ* of the stochastic impedance map due to uncertain reflectance

The estimated values of *μ, μ* + *σ, μ − σ* of stochastic impedance map generated for the PVDF through kriging using linear variogram function is shown in Figure 8 (b), (c) and (d). Table 1 shows the calculation of 10 spatial points selected using Latin hypercube sampling where the index in the table is anamously representing the test point’s location. It can be clearly shown that mean impedance values lie close to the actual values which demonstrates the efficacy and robustness of the proposed framework. The next section will discuss the application of the proposed framework to the fish scale which examines its impedance in the present work.

**Table 1.**
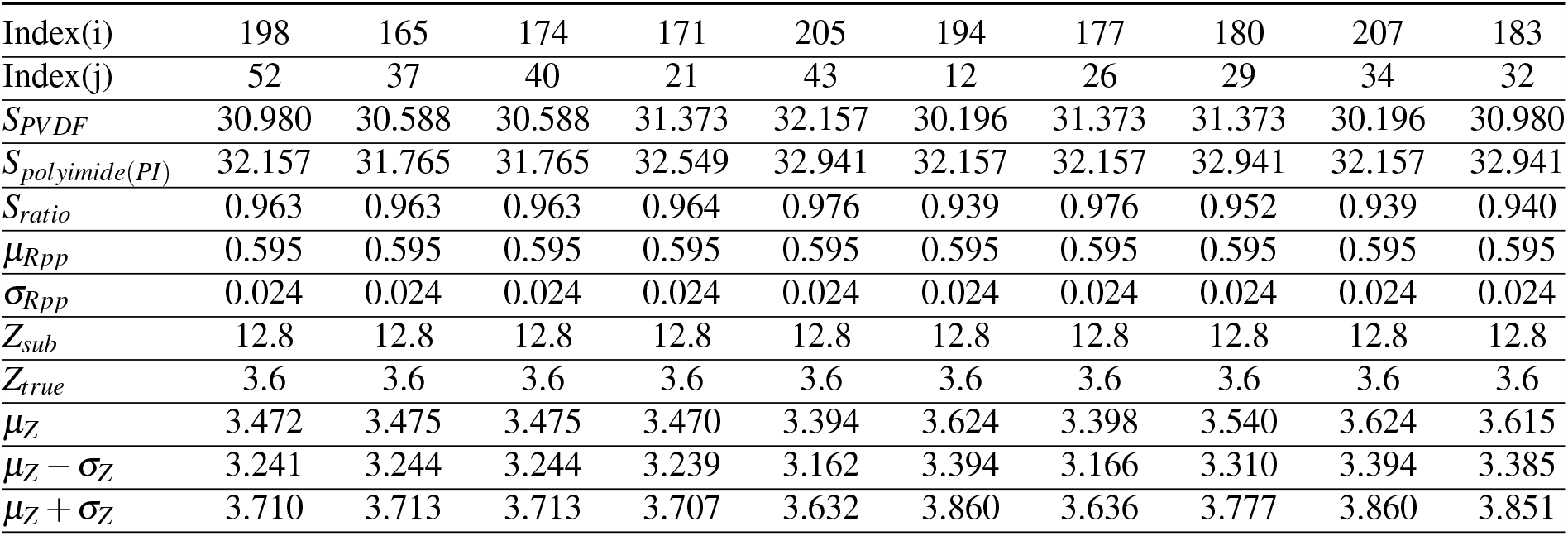
Table 1 represents the corresponding frequency, measured and true impedance of the target and reference specimen and also their uncertainties

|  |  |  |  |  |  |  |  |  |  |  |
| --- | --- | --- | --- | --- | --- | --- | --- | --- | --- | --- |
| Index(i) | 198 | 165 | 174 | 171 | 205 | 194 | 177 | 180 | 207 | 183 |
| Index(j) | 52 | 37 | 40 | 21 | 43 | 12 | 26 | 29 | 34 | 32 |
| $S_{PVDF}$ | 30.980 | 30.588 | 30.588 | 31.373 | 32.157 | 30.196 | 31.373 | 31.373 | 30.196 | 30.980 |
| $S_{polyimide(PI)}$ | 32.157 | 31.765 | 31.765 | 32.549 | 32.941 | 32.157 | 32.157 | 32.941 | 32.157 | 32.941 |
| $S_{ratio}$ | 0.963 | 0.963 | 0.963 | 0.964 | 0.976 | 0.939 | 0.976 | 0.952 | 0.939 | 0.940 |
| $\mu_{Rpp}$ | 0.595 | 0.595 | 0.595 | 0.595 | 0.595 | 0.595 | 0.595 | 0.595 | 0.595 | 0.595 |
| $\sigma_{Rpp}$ | 0.024 | 0.024 | 0.024 | 0.024 | 0.024 | 0.024 | 0.024 | 0.024 | 0.024 | 0.024 |
| $Z_{sub}$ | 12.8 | 12.8 | 12.8 | 12.8 | 12.8 | 12.8 | 12.8 | 12.8 | 12.8 | 12.8 |
| $Z_{true}$ | 3.6 | 3.6 | 3.6 | 3.6 | 3.6 | 3.6 | 3.6 | 3.6 | 3.6 | 3.6 |
| $\mu_Z$ | 3.472 | 3.475 | 3.475 | 3.470 | 3.394 | 3.624 | 3.398 | 3.540 | 3.624 | 3.615 |
| $\mu_Z - \sigma_Z$ | 3.241 | 3.244 | 3.244 | 3.239 | 3.162 | 3.394 | 3.166 | 3.310 | 3.394 | 3.385 |
| $\mu_Z + \sigma_Z$ | 3.710 | 3.713 | 3.713 | 3.707 | 3.632 | 3.860 | 3.636 | 3.777 | 3.860 | 3.851 |

### 5.2 Estimation and development of the impedance map of fish scale

In order to determine the acoustic impedance of the fish scale, we have applied the above-mentioned algorithm to analyze the acoustic response of the fish scale. Here also, the polyimide is used as the base material. The objective now is to isolate the dominant characteristics residing within it through MODWT. To distill the essential characteristics of the signal, it becomes necessary to apply a bandpass filter first, and then decomposition is carried out using MODWT. This decomposition leads to the various time series at different decomposition levels represented by (1), (2), (3), and (4) which are shown in Figure 9. It shows that there is a significant difference in the reflected frequency in both materials even in the decomposed state, the same is clearly visible. This variation in decomposed time series is arising due to wave phenomenon, primarily due to differences in the material impedance and multiple interfaces of reflection. The time series that has maximum energy content is selected as the essential time series and considered as the filtered signal. Figure 10 shows the normalized power of each decomposed time series with respect to others.

**Figure 9.**
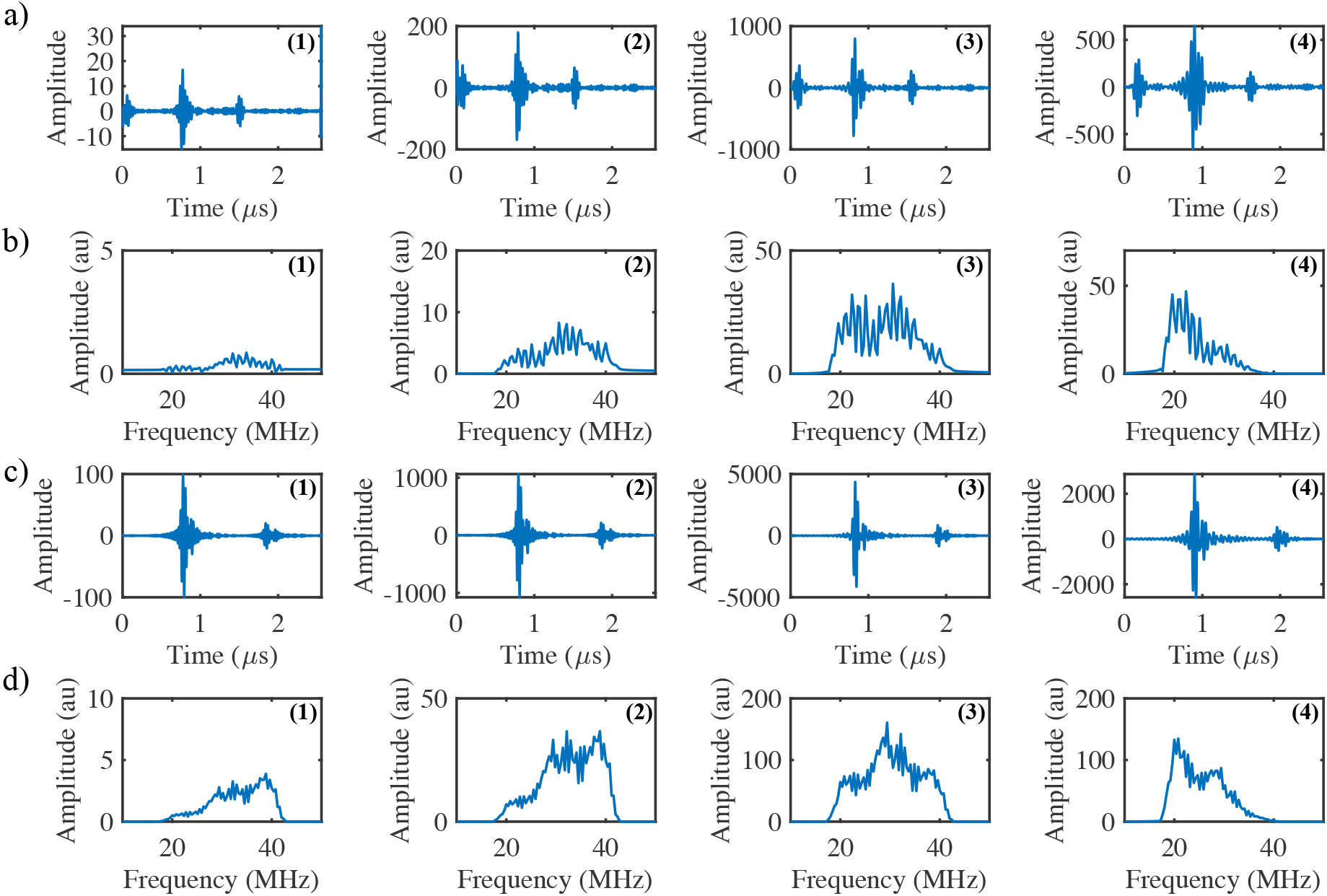
It demonstrates the decomposition of the acoustic response signal into multiple decomposed time series after MODWT in both the time-domain (a), (c) as well as frequency domain (b), (d). Figure (a) and (b) highlights the characteristics of the response signals of fish scale at different decomposition level and figure (c) and (d) highlights the characteristics of the response signal of polyimide (PI) at various decomposition level.

**Figure 10.**
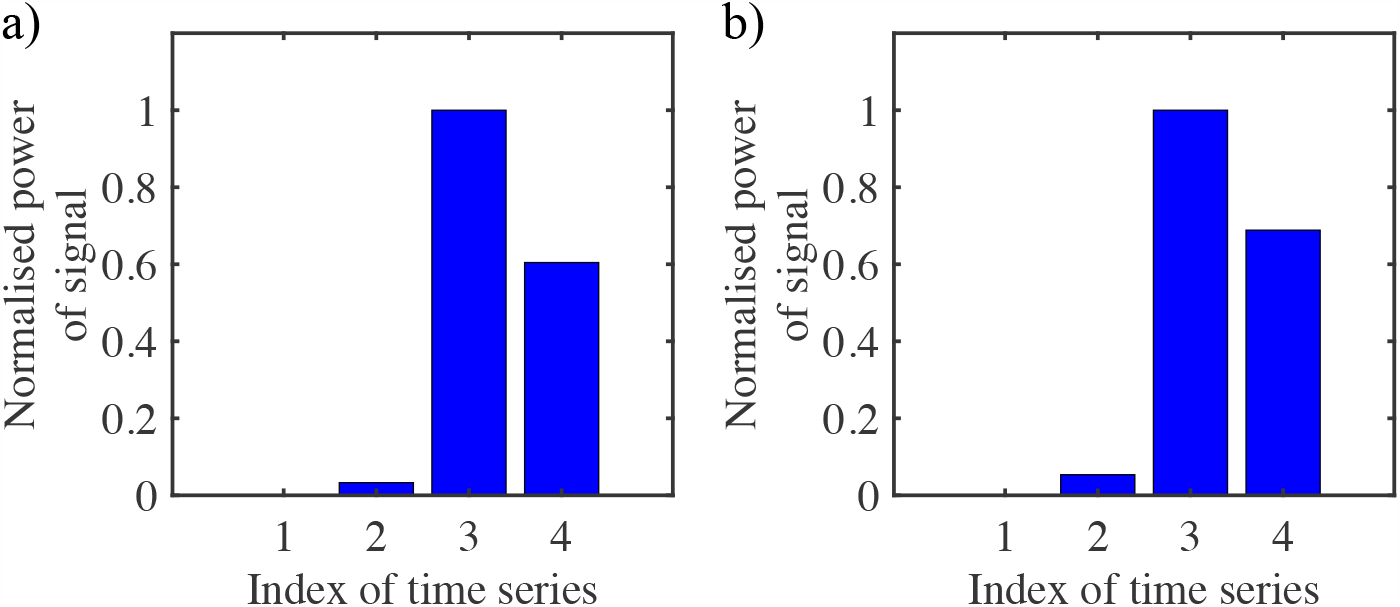
The figures represent the normalized power of each decomposed time series with respect to others for (a) salmon and (b) polyimide (PI) samples, providing the basis for the selection of dominant time series.

As already discussed in the previous case (PVDF and polyimide), the essential characteristics of the signal that are directly correlated to the acoustic properties of fish scales can be extracted from this filtered response forming the fundamental basis for monitoring the bio-mechanical properties through signal processing. However, the changes in the time domain are difficult to interpret and therefore the responses are transformed to the frequency domain where the predominant peak frequency is selected as a representative of the specimen characteristics as presented in Figure 11.

**Figure 11.**
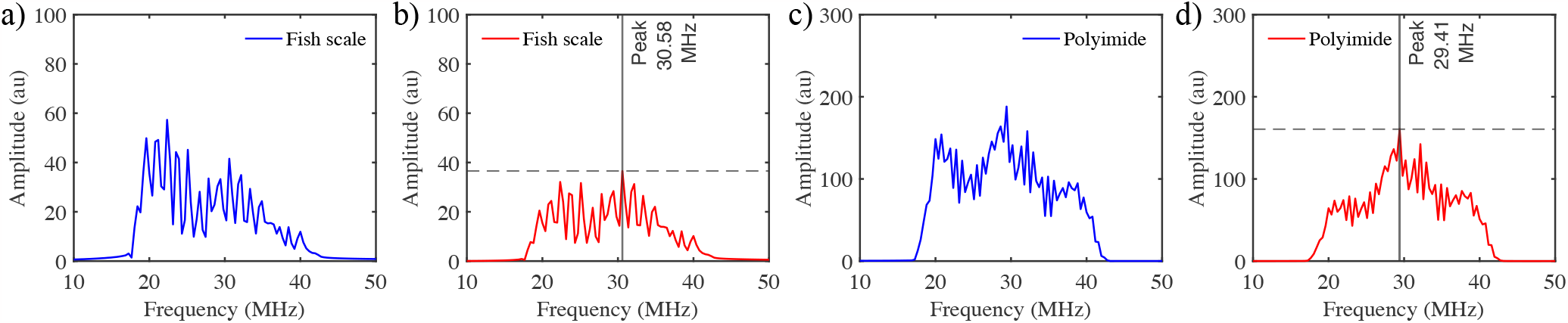
Displayed in the figures are the frequency spectra of the true responses (a) fish scale or scale and (c) polyimide (PI), as well as the wavelet-transformed signals (b) and (d). These representations demonstrate the identification of primary frequencies which is a fundamental step in impedance calculation

Figure 11 (a and c) illustrates the frequency spectra of the true responses of the reference fish scale and target medium polyimide. These obtained frequencies will provide the *S*_*ratio*_ as described in the theory section. It serves as a foundation for impedance calculation, as it reveals the fundamental frequencies correlated to the material that helps us to deduce the reflectance property associated with the medium which ultimately provides the impedance of the fish scale through the procedure described earlier. The frequency spectra of the filtered or wavelet transformed signal are shown in Figure 11 (b and d). The dominant peak from the frequency spectra of the filtered signal is obtained and shown in Figure 11 (b and d). After extraction of dominant frequencies, the stochastic impedance of the fish scale is calculated and thus, the estimated mean value of the acoustic impedance of the fish scale along with uncertainty considered in reflectance is presented in Figure 12 (a) and (b) respectively. The estimated values of *μ −σ, μ* + *σ* of stochastic impedance map generated for the fish scale through kriging using Gaussian variogram is shown in Figure 13 (c) and (d).

**Figure 12.**
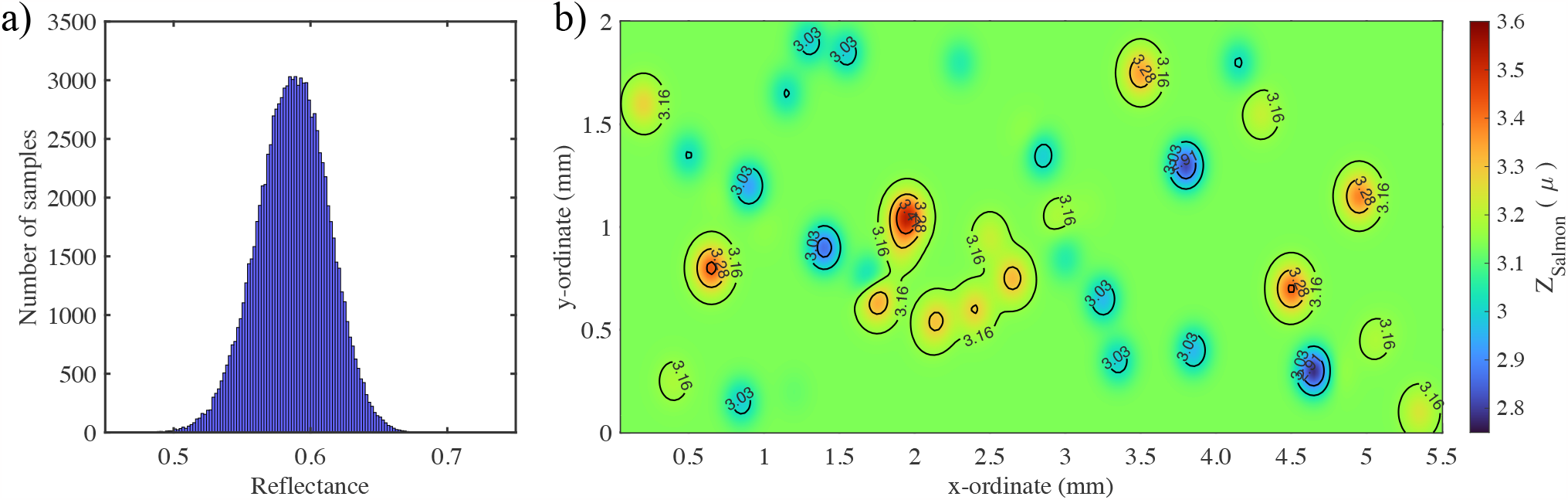
Figure (a) presents the uncertainty considered in the reflectance and Figure (b) illustrates the distribution of estimated mean impedance values, providing a comprehensive view of how impedance varies across the entire domain of interest.

**Figure 13.**
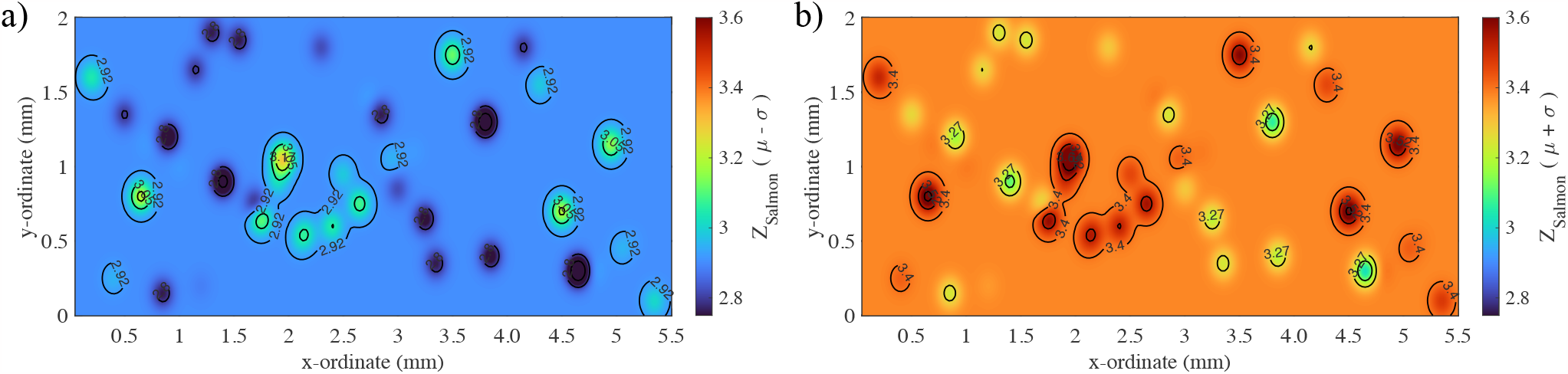
Figure (a) presents the *μ– σ* values of the estimated stochastic impedance maps and Figure (b) illustrates the *μ* + *σ* values of the estimated stochastic impedance maps providing a comprehensive view of how impedance varies across the entire domain of interest with uncertain reflectance.

The figures visually represent the dispersion of estimated impedance values, providing a holistic understanding of how impedance fluctuates throughout the entire area of interest. This representation is indispensable for gaining insights into the impedance characteristics of the fish scale. Further, it can be inferred that the mean impedance values of the considered fish scale lie somewhere between 2.9 to 3.3 Mrayl. Considering, the stochastic reflectance, the values lie between 2.8 to 3.6 Mrayl. These values of impedance can now be used for any further investigations.

## 6 Conclusion

This article introduces a framework designed for the estimation of stochastic impedance through acoustic imaging. This framework is versatile and can be applied to a wide range of samples, including biological specimens such as fish scales as it considers the stochastic reflectance which made its first pioneering application to the fish scale. The framework’s effectiveness is initially established and verified by considering the impedance characteristics of well-known materials. In this context, PVDF is employed as a reference material for calibration to ensure the accuracy of our proposed algorithm. The results confirm that the estimated impedance through the proposed framework lies very nearer to the actual values of impedance for PVDF which shows accuracy exceeding 90%. Additionally, a stochastic impedance map is generated through Kriging using the Gaussian variogram, a technique well-suited for addressing the complexities stemming from spatial variations within the biological specimen. The global analysis of acoustic impedance visualizations provides valuable insights into the functional aspects and complex biomechanical structure of various components within a salmon scale. The methods applied herein may used to study the fish scale regeneration process, as scale loss during delousing strategies and fish skin lesions is of a serious concern in Norwegian aquaculture.

## Acknowledgement

This work was supported by the Research Council of Norway, Cristin project 2061348 ‘VirtualStain’ and project 301401 ‘Nano- and microplastics: Do they impact fish health and welfare’.

## Author contributions statement

A. S., A. H., and S. O. have conceptualized the idea. K. A. designed and performed the experiments. S. O. implemented the algorithm with initial help from A. S. and A. H. Funding was secured by A. H. Formal analysis and experimental validation were performed S.O. S.O. and K.A. also wrote the original draft, and reviewed, and edited the manuscript with support from all co-authors.

## Data availability

The data sets used and/or analyzed during the current study are available from the corresponding author upon reasonable request.

